# Cardiac-confidence coupling is modulated by sensory but not higher-order uncertainty

**DOI:** 10.64898/2026.11.13.765855

**Authors:** Kelly Hoogervorst, Leah Banellis, Siebe Everaerts, Francesco Bavato, Ashley Tyrer, Micah Allen

**Affiliations:** Center of Functionally Integrative Neuroscience, Aarhus University, Aarhus C, 8000, Denmark; Department of Adult Psychiatry and Psychotherapy, University Hospital of Psychiatry Zurich, Zurich, 8032, Switzerland

**Keywords:** Metacognition, Confidence, Cardiac Signals, Interoception, Uncertainty, Emotion Discrimination

## Abstract

Confidence, the subjective sense of certainty accompanying decisions, may be shaped by more than the sensory evidence on which the decision itself was based. Cardiac signals have been linked to confidence, with some evidence that this coupling strengthens when sensory evidence is unreliable. However, sensory evidence can be unreliable in different ways: if sensory input is degraded, predictive cues are misleading, or the environment is too volatile for stable expectations to form. These are distinct computational problems, and whether cardiac-confidence coupling is engaged under all of them has not yet been tested. Similarly, whether this coupling predicts individual differences in metacognitive efficiency remains unknown. To address this, we recorded cardiac activity from 167 participants performing a face-emotion discrimination task that orthogonally manipulated sensory precision, cue validity, and block volatility, with trial-wise confidence ratings. Pre-decisional heart rate deceleration was more pronounced under low than high sensory precision. Within this window, trial-level heart rate positively predicted confidence, independently of accuracy and precision. This cardiac-confidence coupling was stronger when sensory precision was low, but cue validity and block volatility did not modulate the coupling. Neither individual resting heart rate variability nor coupling strength predicted metacognitive efficiency. Together, these findings indicate that cardiac-confidence coupling is modulated by sensory uncertainty but not by higher-order uncertainty, consistent with a within-trial interoceptive weighting mechanism rather than a stable individual-differences trait.

## Introduction

Decisions are accompanied by confidence, a subjective estimate of the probability of being correct. This estimate is a second-order computation, drawing on sources of information that may be partly independent of first-order processes (Fleming, 2024). Confidence has been shown to calibrate learning, govern whether a choice gets revised, and modulate the weight of current evidence relative to new input (Pouget et al., 2016; Yeung & Summerfield, 2012). Alterations in metacognition have been associated with psychiatric conditions, such as reduced confidence under anxious-depressive symptoms (Rouault et al., 2018; Seow et al., 2021). Understanding its contributions is therefore essential for understanding how confidence is computed, and when and why it fails.

A growing body of evidence points to physiological signals as one such input. Cardiac afferents reach the brain via vagal pathways through the nucleus of the solitary tract, projecting to the insular cortex, anterior cingulate, and prefrontal regions implicated in metacognitive computation (Critchley et al., 2004; Fleming & Dolan, 2012; Park & Blanke, 2019). Previous research suggests that these signals are not just treated as background noise when they reach the brain, but that they are integrated into our higher-order cognitive processes (Critchley & Harrison, 2013; Seth, 2013). In particular, previous research has found that heart rate typically decelerates in the moments before a decision or stimulus, a response commonly referred to as the cardiac orienting response (Bradley, 2009; Graham & Clifton, 1966; Lacey & Lacey, 1978). This orienting response is hypothesised to sharpen processing of sensory evidence by boosting the signal-to-noise ratio (SNR) of incoming information (Bradley, 2009; Porges, 2007). One proposed mechanism is that deceleration suppresses the afferent baroreceptor signals, thereby reducing baroreceptor-mediated inhibition of cortical processing, attenuating interoceptive interference and freeing up resources for external input (Lacey & Lacey, 1978; Skora et al., 2022).

Beyond their role in sharpening sensory processing, cardiac signals may also contribute to higher-order inference. From a predictive processing perspective, multiple evidence sources are weighted by their relative precision (Feldman & Friston, 2010), so when sensory evidence is weak, internal information is expected to increase in relative weight (Seth & Friston, 2016). Accordingly, if confidence integrates cardiac information, it should track cardiac signals more closely when sensory evidence is weak (Allen et al., 2022). Consistent with this, cardiac activity has been shown to influence perceptual decisions and confidence, with stronger effects when sensory evidence is unreliable (Allen et al., 2016; Garfinkel et al., 2014). However, there are multiple levels at which our environment can be manipulated and uncertainty introduced.

Uncertainty can arise because sensory input is noisy, because predictive cues are misleading on a given trial, or because the environment is too volatile for stable expectations to form across trials. These are different computational problems operating at different levels of inference: the sensory likelihood, the learned prior, and the rate at which that prior should update (Mathys et al., 2014). Whether interoceptive contributions to confidence are engaged under all of these instances, or selectively when the sensory evidence itself is degraded, has not been directly tested. At the individual level, a separate question arises. If cardiac signals carry information relevant to metacognition, then individuals who integrate this signal more strongly should have access to additional information and, in principle, display higher metacognitive efficiency. Similarly, resting heart rate variability (HRV), a marker of autonomic flexibility linked to top-down regulation and adaptive belief updating (Forte & Casagrande, 2025; Thayer et al., 2009), could facilitate the dynamic weighting of interoceptive and exteroceptive evidence. Whether either of these cardiac metrics predicts metacognitive efficiency remains untested.

The present study addressed these questions using the confidence weighting task (CWT), a novel face-emotion discrimination paradigm that orthogonally manipulates sensory precision, cue validity, and blockwise volatility with simultaneous electrocardiography (ECG) recordings. We aimed to establish, first, whether trial-level heart rate predicts confidence after controlling for accuracy, to separate cardiac-confidence coupling from task performance; second, whether this coupling strengthens when sensory evidence is weak, and whether this modulation extends to higher-order sources of uncertainty; and third, whether individual differences in resting vagal tone or cardiac-confidence coupling strength predict metacognitive efficiency.

## Methods

### Participants

Participants were recruited within the second cohort of the Visceral Mind Project (VMP2), a large-scale data-collection project conducted at the Center of Functionally Integrative Neuroscience (CFIN) at Aarhus University. The VMP involved multiple behavioural and neuroimaging sessions; the present study used data from the CWT with concurrent ECG recordings and additional photoplethysmography (PPG) recordings from the resting-state functional magnetic resonance imaging (MRI) session. Inclusion criteria required normal or corrected-to-normal vision, fluency in Danish or English, as well as compatibility with MRI requirements (no metal implants, no claustrophobia, not pregnant or breastfeeding), and no history of neurological diagnoses or use of medications other than over-the-counter antihistamines or hormonal contraceptives. All participants provided written informed consent and were remunerated for their participation. The Regional Ethics Committee of Region Midtjylland granted ethical approval (78118), and the study was conducted in accordance with the Declaration of Helsinki.

Of the 285 participants who completed the CWT task (191 women, 94 men; mean age 24.6, *SD* = 4.5, age range = 18-52), 19 did not have physiological data collected and were excluded, leaving 266 with both behavioural and cardiac recordings. A further total of 99 participants were excluded through a staged quality-control pipeline. First, participants were excluded based on the quality and completeness of physiological recordings (*n* = 34): poor or unusable signal quality (*n =* 6), incomplete recording shorter than 25 minutes (*n* = 6), failure of photodiode-based trigger alignment (*n* = 7), usable photodiode triggers on fewer than 80% of trials (*n* = 2), or more than 5% of task-period beats flagged as artefacts (*n* = 13). Second, participants were excluded based on task performance and confidence-rating quality (*n* = 65 unique): fewer than 80% of trials retained after removing reaction-times below 50 ms (*n* = 4), mean accuracy below 60% on hard (low-precision) trials or below 80% on easy (high-precision) trials (*n* = 25); within-subject confidence standard deviation below 10 on a 0-100 scale (*n* = 6); mean confidence below 10 or above 90 (*n* = 27); or more than 70% of confidence responses at the scale endpoints of 0 or 100 (*n* = 30). These were a priori data-quality filters chosen to retain participants with the dynamic range in accuracy and confidence needed for reliable metacognitive estimation. The accuracy bounds exclude performance at or near chance; the confidence criteria exclude ratings with no variability or clipped at the scale extremes. Confidence was only assessed on trials where the participant had made a perceptual response. The final analytic sample comprised 167 participants (110 women, 57 men; mean age 24.6 years, *SD* = 4.2, age range = 18-45). Participants self-reported gender. No sex- or gender-based analyses were conducted. Resting-state HRV was unavailable for participants who did not complete the 14-minute fMRI session (*n* = 7), yielding *N* = 160 for analyses involving this measure. All other analyses used the full sample of 167.

### Procedure

Prior to the CWT, participants completed a Face Affect Discrimination (FAD) thresholding task to establish their individual psychometric threshold for distinguishing between angry and happy facial expressions (∼5 min). Face stimuli were drawn from a morph continuum ranging from fully angry (morph level 0) to fully happy (morph level 200) to produce a valence gradient (see *Stimuli*). Sensory precision refers to the clarity of the emotional signal: extreme morphs (0 or 200) are high-precision and intermediate morphs are low-precision. Accordingly, high- and low-precision stimuli were drawn from the extreme and intermediate morphs, respectively (see *Task design*). The task used a Psi adaptive procedure based on Kontsevich & Tyler (1999), implemented in the Palamedes toolbox (Prins, 2013), fitting a Weibull psychometric function over 60 trials to jointly estimate the threshold (α, the morph level corresponding to the point of subjective equality) and slope (β) parameters. The 80%-correct morph levels used as low-precision stimuli in the main task were then derived from this fitted function per emotion. Each thresholding trial consisted of a face stimulus presentation (1.0 s), an emotion decision (up to 1.0 s), and a variable inter-trial interval (ITI; 1.0-2.0 s). Following the thresholding task, participants received written instructions and completed a 12-trial practice session (2 instructional trials plus 10 practice trials).

### Task design

The CWT adapted the probabilistic learning task of Lawson et al. (2021), replacing their face-versus-house discrimination with added white visual noise with a happy-angry morph continuum as the sensory precision manipulation. In the CWT, each trial proceeded as follows: a cue image was presented (0.5 s), followed by a variable inter-stimulus interval (ISI; 2.0-3.0 s) with a central fixation cross, and then a face stimulus (0.5 s). Participants indicated whether the face appeared more angry or happy via button press (max 2.0 s), and subsequently provided a confidence rating on a continuous visual analogue scale (VAS; 0-100, anchored from “Guess” to “Certain”) using a mouse slider (max 3.0 s). Trials were separated by a variable ITI (1.0-2.0 s; Figure 1A).

**Figure 1:**
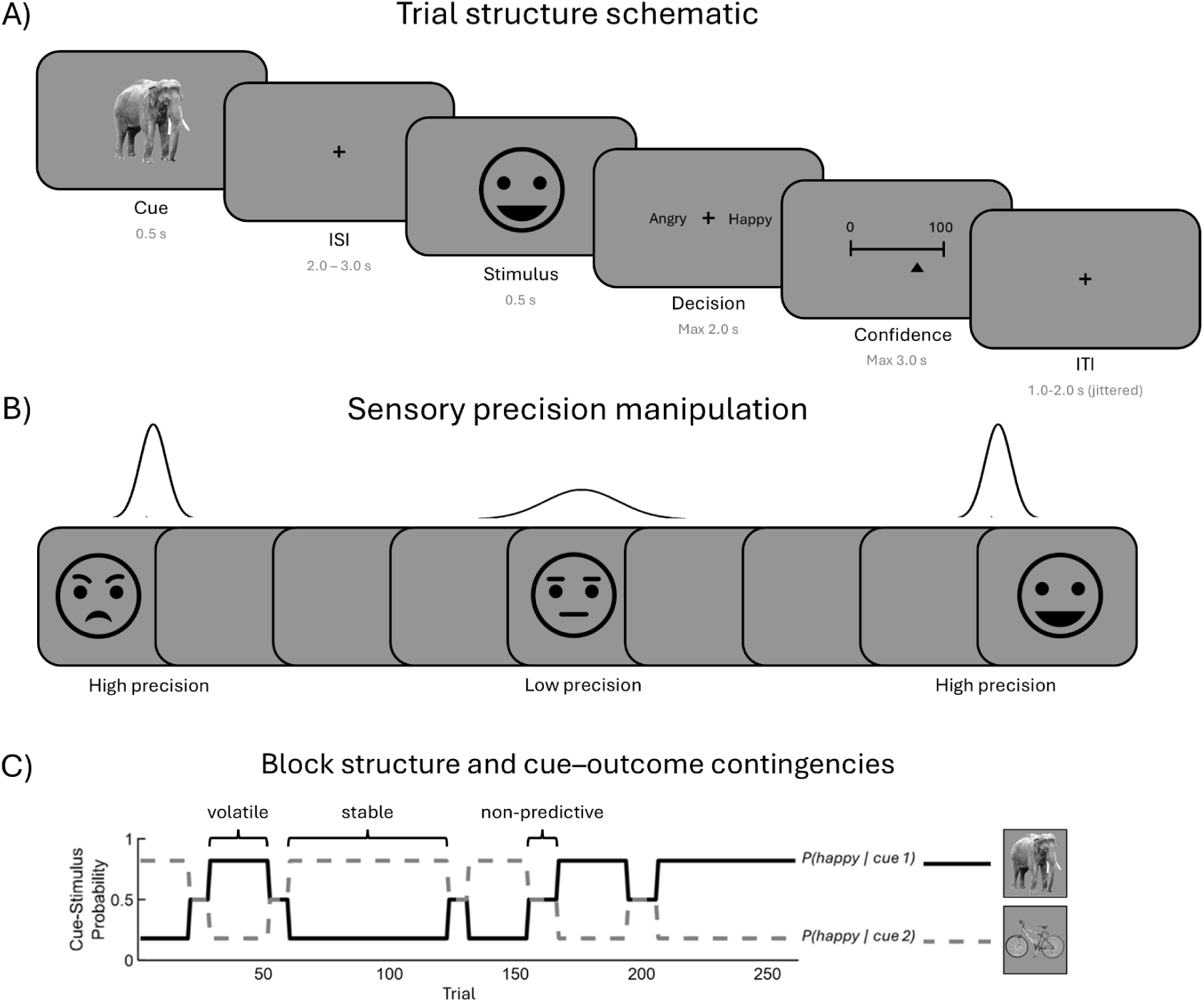
Confidence Weighting Task (CWT) schematic. A) Trial structure. Each trial began with a cue image (0.5 s), followed by a fixation interval (2.0-3.0 s), a face stimulus (0.5 s), an emotion decision (angry/happy; max 2.0 s), a confidence rating on a continuous visual analogue scale (0-100; max 3.0 s), and a jittered inter-trial interval (1.0-2.0 s). B) Sensory precision manipulation. Face stimuli were drawn from a morph continuum ranging from fully angry to fully happy. High-precision stimuli (extremes of the continuum) were easy to discriminate; low-precision stimuli (near the centre) were individually calibrated to each participant’s threshold. C) Block structure and cue-outcome contingencies. Solid and dashed lines show the probability of a happy face given each cue across the session. In stable blocks, one cue predicted happy faces and the other predicted angry faces (82% validity). In volatile blocks (short segments), contingencies shifted quickly. In non-predictive blocks, both cues were equally associated with each emotion (50%). Brackets label the three block types. Face icons in (A) and (B) are schematic illustrations only; the actual face stimuli were morphed composite images derived from the Karolinska Directed Emotional Faces (KDEF) database (Lundqvist et al., 1998).

The task orthogonally manipulated three sources of uncertainty. Sensory precision reflected how clearly the face expressed an emotion; cue validity reflected whether the cue’s prediction matched the face emotion on a given trial; and volatility reflected how long the block’s cue-emotion contingencies held before changing. Sensory precision was varied at two levels per emotion. High-precision stimuli used fixed morph endpoints (morph level 0, fully angry; morph level 200, fully happy; easy-to-discriminate; Figure 1B). Low-precision stimuli were individually selected for each emotion to target ∼80% correct discrimination on the FAD, a standard psychophysical operating point where discrimination is above chance but error-prone. Cue validity was manipulated via two cue images (an elephant and a bicycle) that predicted the upcoming face emotion. In predictive blocks, one cue validly predicted the face emotion on 82% of trials and was invalid on the other 18%. Volatility was manipulated across blocks: stable blocks were relatively longer (∼60 trials) with fixed cue-outcome contingencies maintained throughout, whereas volatile blocks were shorter (∼24 trials), leaving only a brief period to learn the cue-outcome contingency (Figure 1C). Non-predictive blocks (∼10 trials each), in which cues carried no information about the upcoming face emotion, were intermixed throughout the session. The full task comprised 11 blocks (2 long + 4 short + 5 non-predictive) totalling 264 main trials, with two self-paced breaks placed at approximately one-third and two-thirds of the way through the session.

Approximately every 12 standard trials, a prediction trial was inserted (22 total), in which the preceding cue was re-presented, and participants indicated which face emotion they expected that cue to predict. Prediction trials were not included in subsequent analyses. Visual synchronisation markers were embedded in a corner of the display via a photodiode to enable precise temporal alignment between behavioural and physiological data.

### Stimuli

Face stimuli were drawn from a 201-step morph continuum generated from two gender-specific KDEF composite anchor faces (one male, one female), each averaged from 9 KDEF identities per emotion category. The composites and morph continuum were developed by Brændholt et al. (2025) based on the Karolinska Directed Emotional Faces (KDEF) database (Lundqvist et al., 1998), ranging from 100% angry (morph level 0) to 100% happy (morph level 200) in 201 discrete steps. Stimuli were balanced across gender (50% male, 50% female). High-precision stimuli corresponded to morph levels 0 (fully angry) and 200 (fully happy). Low-precision stimuli corresponded to each participant’s individually estimated morph levels calibrated to a predicted ∼80% correct discrimination accuracy per emotion. All face stimuli were displayed at 4° of visual angle at a viewing distance of 56 cm. Cue stimuli were drawn from the Bank of Standardized Stimuli image set (Brodeur et al., 2014), and consisted of an elephant and a bicycle, selected for minimal inherent valence associations, presented at the same visual angle as the face stimuli.

### ECG acquisition

Cardiac activity was recorded continuously throughout the task using a bipolar surface ECG configuration (Ag/AgCl electrodes placed below the right and left collarbones, with a ground electrode on the left rib curvature), amplified and digitised at 1000 Hz using a biosignalsplux system (PLUX Wireless Biosignals, Lisbon, Portugal). The photodiode signal was recorded on a parallel channel to enable precise alignment of cardiac data with behavioural timestamps. In addition, PPG was recorded during a separate resting-state fMRI session for the computation of resting heart rate variability (see *Cardiac metrics*).

### ECG preprocessing

Raw ECG signals were preprocessed using a custom Python pipeline. To remove baseline drift and high-frequency artefacts, signals were band-pass filtered using a 4th-order zero-phase Butterworth filter (1-40 Hz), followed by a 50 Hz notch filter (Q = 50) to attenuate electrical mains interference. Filtered signals were downsampled to 300 Hz using polyphase resampling. Signal quality was assessed using seven automated checks covering flatline detection, signal clipping, signal-to-noise ratio (SNR; threshold ≥ 10 dB), R-peak detection rate, heart rate plausibility, QRS amplitude consistency, and baseline drift. These checks assessed overall recording quality, and each check produced a binary pass/fail result and a weighted quality score (0-1) was computed across checks. Signal quality was additionally assessed by visual inspection of the ECG trace, the detected R-peak sequence, and the resulting RR intervals.

Cardiac data were aligned to behavioural timestamps using photodiode-based trigger detection. The trigger sequence was matched against the expected cue-face-decision pattern from the first 15 behavioural trials using pattern matching, with the alignment offset defined as the difference between the first detected trigger time and the first behavioural cue onset. Alignment was accepted only if at least 85% of pattern triggers matched; recordings below that threshold were excluded. Alignment quality was then further verified by computing the inter-trial interval (ITI) correlation between photodiode triggers and behavioural timestamps, and the mean timing error across the first 50 trials. Recordings with ITI correlation < 0.98 or mean timing error > 100 ms were flagged for manual inspection of the photodiode signal.

R-peaks were detected using NeuroKit2’s default detector (Makowski et al., 2021). To supplement detection in the first 3 seconds of the recording, an additional threshold-based detector was applied to the first 900 samples. Detected peaks were refined by shifting each detection to the true local signal maximum within a ±37 ms window. Peaks were removed when both adjacent RR intervals were physiologically implausible (< 430 ms or > 1500 ms, corresponding to > 140 bpm or < 40 bpm); flagged intervals themselves were excluded from HRV and from the HR spline. Artefacts were identified using a local variability method: any RR interval deviating by more than 20% from the mean of the surrounding 10 intervals was flagged. Flagged intervals were excluded from downstream analyses; detected R-peak timestamps were passed through unchanged. Preprocessed cardiac data were trimmed to the task window (5 seconds before the first cue onset to 15 seconds after the final cue onset) and peaks falling within rest breaks (inter-trial gaps > 10 seconds) were flagged and excluded from subsequent analyses. Per-trial alignment was refined using the photodiode triggers recorded on the parallel channel. For each trial, the residual clock drift between the behavioural log and the photodiode-detected cue onset was measured. A trial’s drift estimate was accepted when the cue-trigger and face-trigger drifts agreed within 50 ms, and the estimate did not deviate by more than 100 ms from the median of the five neighbouring trials in each direction. Rejected or missing drift estimates were linearly interpolated from accepted trials.

### Cardiac metrics

Instantaneous heart rate (HR) during the task was estimated by fitting a gap-aware cubic spline through the timestamps and corresponding HR values of artefact-free beats. The spline was only evaluated at time points where the two flanking clean beats were no more than 3 s apart (twice the longest accepted RR interval); inside longer gaps (artefact runs and rest breaks), HR was set to missing. For trial-locked analyses, HR was sampled in 500 ms bins and baseline-corrected by subtracting the mean HR in the 1 s preceding cue onset. Baseline HR was set to missing when fewer than 50% of samples in the baseline window were supported by the spline.

Resting-state heart rate variability (HRV), indexed by the root mean square of successive differences (RMSSD), was derived from photoplethysmography (PPG) recorded during a resting-state fMRI scan in a separate session within the VMP2; the fMRI data are not reported here. PPG-derived RMSSD closely approximates ECG-derived RMSSD at rest (Schäfer & Vagedes, 2013); we use the term HRV throughout. Systolic peaks were identified using the systole package (Legrand & Allen, 2022) with manual visual inspection and correction, following the protocol described in Banellis et al. (2026). RMSSD was log-transformed, and values more than 3 SD from the mean of the log-transformed distribution were treated as outliers and set to missing (Task Force, 1996). No participants exceeded this threshold (maximum |*z*| = 2.67; range: 10.6-233.7 ms).

### Data analysis

The analyses addressed three questions: what modulates the cardiac response itself, what modulates the strength of cardiac-confidence coupling, and whether individual differences in cardiac measures predict metacognitive efficiency. Below, we first describe the behavioural models, then the time-resolved cardiac and coupling analyses, and finally the individual-differences analyses.

Behavioural analyses used Bayesian multilevel models fitted with brms (version 2.23.0; Bürkner, 2017) using the cmdstanr backend. Perceptual sensitivity (*d*′) was estimated using a probit signal-detection model treating happy as signal and angry as noise, with precision as a fixed effect and participant as a random effect. Reaction times were modelled with a lognormal family, and confidence ratings (0-100) with an ordered beta family appropriate for bounded continuous data. Both included fixed effects of accuracy, precision, and their interaction, with random intercepts and slopes for accuracy, precision, and their interaction by participant. Separate ordered-beta models were also fitted for confidence as a function of cue validity (low-precision, stable blocks) and block volatility (low-precision trials, non-predictive blocks excluded), each with the condition as a fixed effect and random intercepts and slopes by participant. Weakly informative priors were placed on all fixed effects: normal(0, 3) for the d′ model, normal(0, 1) for the RT model, and a coefficient prior SD of 2.5 with ordbetareg defaults for the four ordered-beta confidence models.

Models were fitted with 4 chains (warmup = half). Four models required 4,000 iterations per chain and two required 2,000; adapt_delta was raised from 0.80 to 0.95 for the confidence × precision and confidence × cue validity models after initial fits produced divergent transitions. Convergence was assessed via R-hat (< 1.01), effective sample size (ESS), and the absence of divergent transitions. All six models produced zero divergent transitions, and five met R-hat < 1.01 across all parameters. The confidence-by-precision model (Figure 2D) reached R-hat 1.014-1.016 on two population-level coefficients, with bulk ESS above 600 and estimates stable to within 0.1 scale points across a threefold increase in sampling. In the d’ model (Figure 2A), maximum R-hat of 1.028 was confined to one weakly identified random-effect correlation; all population-level coefficients passed (R-hat ≤ 1.003). Results are reported as posterior medians with 95% credible intervals (Crl) and the probability of direction (*pd*). Frequentist mixed models (lme4/lmerTest; Bates et al., 2015; Kuznetsova et al., 2017) and paired Wilcoxon signed-rank tests are reported as robustness checks. Because confidence ratings clustered at ceiling under high sensory precision (64.0% of high-precision trials at the scale maximum, versus 11.6% of low-precision trials), the cue validity and block volatility analyses were restricted to low-precision trials, where the confidence distribution retained its full range. Non-predictive blocks, in which cues carried no information about the upcoming face emotion, were excluded from the block volatility analysis, which compared stable with volatile predictive blocks.

**Figure 2:**
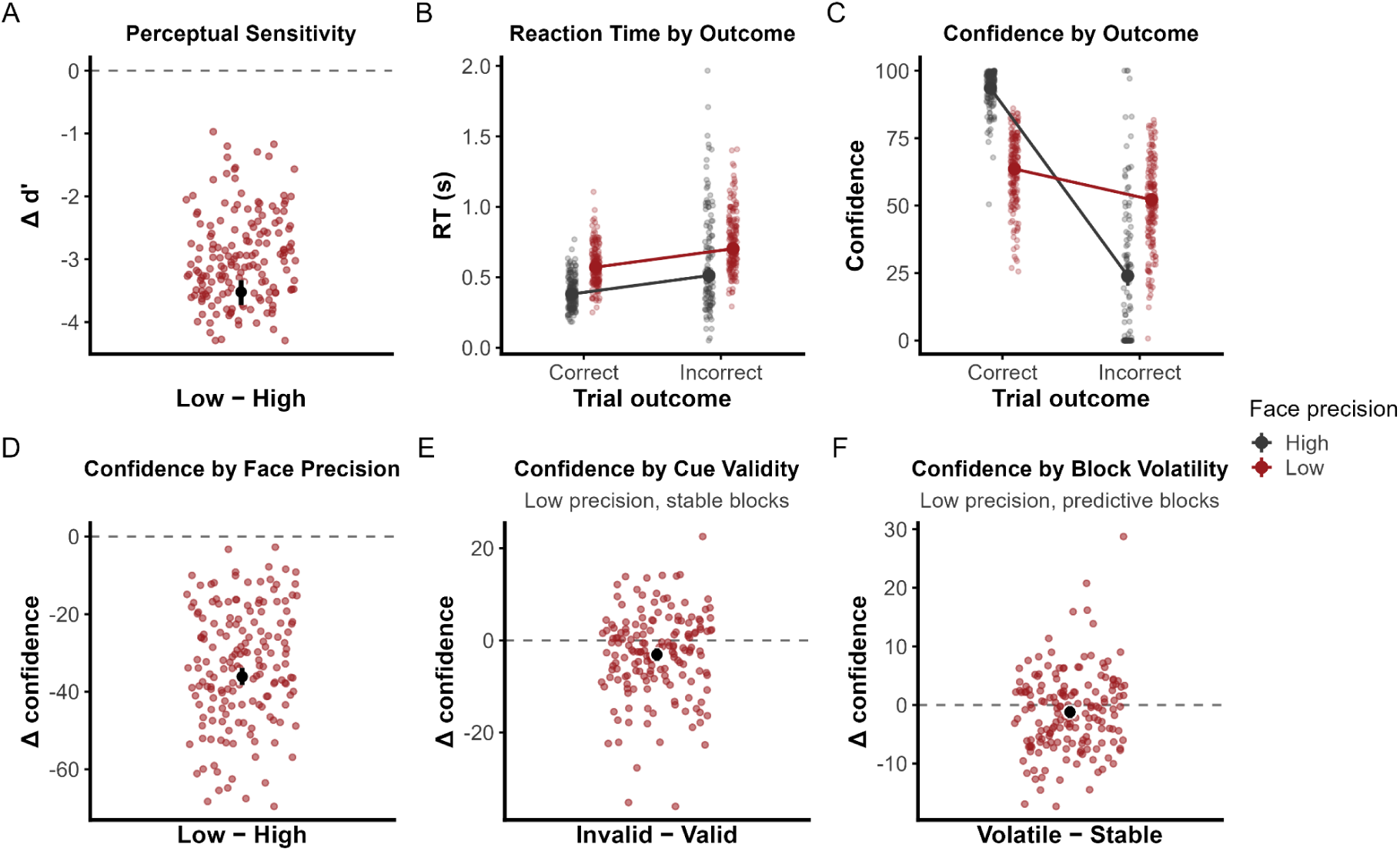
Behavioural performance and confidence across uncertainty manipulations. A) Participant-level difference in *d*′ (low − high precision). B) Reaction time by trial outcome and precision; faded points show participant medians, solid symbols and connecting lines show population-level posterior medians. C) Confidence by trial outcome and precision; faded points show participant means, solid symbols show marginal posterior means. The high-precision incorrect cell contains 253 trials from 110 of 167 participants. D) Participant-level difference in overall confidence (low − high precision). E) Participant-level difference in confidence by cue validity (invalid − valid; low-precision, stable blocks). F) Participant-level difference in confidence by block volatility (volatile − stable; low-precision trials). In A and D- F, each point is one participant’s difference score; the large dot and error bar show the posterior median and 95% CrI. *N* = 167.

To characterise how heart rate evolved during a trial, we computed trial-locked heart rate time courses for each participant. HR was sampled in 500 ms bins and baseline-corrected by subtracting the mean heart rate in the 1 s preceding cue onset. Epoch windows and locking events are specified per analysis below. For all time-resolved cardiac analyses, a linear mixed-effects model (lme4/lmerTest; Bates et al., 2015; Kuznetsova et al., 2017) was fitted independently at each time bin, yielding one t-statistic per bin for the effect of interest. Statistical inference on the resulting t-statistic time series used cluster-based permutation testing (Maris & Oostenveld, 2007) with 1,000 permutations. The cluster-forming threshold was set at the 97.5th percentile of the t-distribution with df = n − 1, where n was the median number of participants contributing to a time bin. Adjacent bins exceeding this threshold with the same sign were grouped into clusters, and cluster mass was computed as the sum of their t-statistics. The cluster-level *p*-value was computed as *p* = (k + 1)/(n_perm + 1), where k is the number of permutations yielding a cluster mass at least as extreme. To generate the null distribution, the sign of each participant’s data was randomly flipped, and the per-bin models were re-fitted. Tests were two-tailed at α = .05 unless otherwise noted. The grand-mean baseline-corrected heart rate was tested against zero to locate periods of significant deviation from baseline. To test whether the deceleration differed by precision, we fitted HR ∼ precision + (1|sID) at each bin, with high precision as the reference level. Equivalent analyses tested whether the deceleration differed by cue validity (low-precision trials, stable blocks) or block volatility (low-precision trials, non-predictive blocks excluded). As a supplementary analysis, we tested whether face-locked heart rate differed by stimulus emotion (happy vs angry) and its interaction with precision (HR ∼ emotion × precision + (1|sID)).

We then ran five cluster-based analyses testing whether trial-level heart rate predicted behaviour. Heart rate and confidence were z-scored within participant across all conditions, so that condition-specific means were preserved. First, as a control, we tested whether heart rate predicted accuracy (accuracy ∼ HR_z + confidence_z + precision + (1|sID)), time-locked to face onset (−3.0 to 3.5 s; two-tailed); because accuracy is binary, we confirmed this result with a logistic mixed model as a robustness check. Second, we tested whether heart rate predicted confidence (confidence_z ∼ HR_z + accuracy + precision + (1|sID)), time-locked to rating scale onset (−4.0 to 3.0 s; one-sided positive). Third, we tested whether this coupling was modulated by sensory precision (confidence_z ∼ HR_z × precision + cue_validity + accuracy + (1|sID); one-sided positive, high precision as reference). Fourth and fifth, we tested equivalent interactions for cue validity (HR_z × cue_validity; two-tailed) and block volatility (HR_z × BlockVolatility; two-tailed); both controlled for accuracy with random intercepts for participants. The validity analysis was restricted to low-precision trials in stable blocks; the volatility analysis to low-precision trials with non-predictive blocks excluded. To assess whether the cardiac-confidence coupling could be explained by shared trial difficulty, we repeated the confidence analysis within low-precision correct trials, adding log-transformed RT as a covariate and nesting participants within face outcome (confidence_z ∼ HR_z + logRT_z + (1|sID/face_outcome)).

Finally, we tested whether individual differences in resting vagal tone or cardiac-confidence coupling predicted metacognitive efficiency. Metacognitive efficiency was indexed by the M-ratio (meta-*d*′/*d*′) estimated using the hierarchical HMeta-d model (Fleming, 2017), fitted with 3 chains of 10,000 samples after 1,000 burn-in. Confidence ratings were binned into 4 equal-width bins within low-precision trials. Each cardiac predictor (z-scored across participants) was entered as a covariate in a separate hierarchical regression on log(M-ratio). HR-confidence coupling strength was extracted per participant by fitting OLS regressions (confidence_z ∼ HR_z + accuracy) at each time bin within the −0.5 to 1.0 s window relative to rating scale onset, within low-precision trials, and averaging the HR_z coefficient across bins. To quantify evidence for null results, Bayes factors (BF₀₁) were obtained via the Savage-Dickey density ratio for the individual-differences analyses (Figure 5), under N(0, 1), N(0, 0.5), and N(0, 0.25) priors on the standardised regression coefficient. Window-based BF₀₁ for Figure 4 (A, D, E) averaged per-bin β within an a priori window (−0.5 to 1.0 s for coupling; 0.0 to 3.5 s post-face for HR → accuracy) and tested the mean against zero (JZS one-sample t-test; BayesFactor package; Morey et al., 2026). These quantify evidence for the null within the window, not across the full time course; prior-width sensitivity is reported.

Analysis code and anonymised pre-processed data supporting the findings of this study will be made publicly available upon publication.

## Results

### Higher perceptual sensitivity and confidence for high-precision stimuli

Overall task performance differed significantly between face precision conditions. Perceptual sensitivity (*d*′) was substantially higher in the high-precision condition (4.90 [4.72, 5.09]) than in the low-precision condition (1.37 [1.29, 1.46]; difference = −3.52 [−3.73, −3.33], *pd* = 1.000; Figure 2A), confirming that the precision manipulation successfully modulated perceptual discriminability.

Participants responded more slowly on incorrect than correct trials and more slowly on low- than high-precision trials. The accuracy effect was present in both precision conditions (high: 0.130 [0.077, 0.187] s, *pd* = 1.000; low: 0.131 [0.116, 0.146] s, *pd* = 1.000), but there was no evidence for an interaction on the absolute scale (0.001 [−0.057, 0.055], pd = 0.511). A frequentist check on log-transformed RT detected a significant multiplicative interaction (F(1, 23340) = 12.77, p < .001), reflecting the proportionally larger slow-down relative to the faster high-precision baseline (0.46 s vs 0.73 s). Because the absolute accuracy-related slow-down (∼130 ms) did not differ between conditions, we treat the absolute-scale result as primary. Cue-validity priming on RT was present at both precision levels and did not differ between them, confirming that participants utilised cue-stimulus information regardless of stimulus precision. RT and accuracy across all three manipulations are reported in Supplementary Table 1.

Confidence was higher on correct than incorrect trials in both precision conditions. Under high precision, confidence was near ceiling on correct trials (97.43 [96.61, 98.11]) and dropped sharply on incorrect trials (12.46 [7.40, 18.70]), resulting in a confidence-accuracy gap of −84.95 [−90.15, −78.65] (*pd* = 1.000). Under low precision, the gap was much smaller: correct 64.19 [62.17, 66.20], incorrect 52.03 [49.40, 54.62], difference −12.15 [−13.63, −10.77] (*pd* = 1.000). The accuracy × precision interaction was large and decisive (72.75 [66.38, 78.08], *pd* = 1.000; Figure 2C).

Overall confidence was substantially higher in the high-precision condition (97.05 [96.11, 97.79]) than in the low-precision condition (60.93 [58.91, 62.95]; difference = −36.10 [−38.25, −33.87], *pd* = 1.000; Figure 2D), indicating that participants were not only more accurate but also more confident when faces were easier to discriminate. Focusing on low-precision trials in stable blocks, confidence was higher on valid-cue trials (61.31 [59.23, 63.38]) than on invalid-cue trials (58.29 [55.98, 60.48]; difference = −3.09 [−4.45, −1.75], *pd* = 1.000; Figure 2E), suggesting that when perceptual information was ambiguous, anticipatory cues guided confidence. Confidence was slightly higher in stable than volatile blocks within low-precision trials (stable: 60.64 [58.52, 62.65]; volatile: 59.42 [57.13, 61.66]; difference = −1.21 [−2.25, −0.19], *pd* = 0.988; Figure 2F). This suggests that greater environmental uncertainty modestly reduced subjective confidence, independent of perceptual difficulty.

### The cardiac orienting response shows sensitivity to sensory precision

During the task, participants’ mean heart rate was 76.6 BPM (*SD* = 10.3; Figure 3A), and mean resting-state HRV (RMSSD) was 54.8 ms (*SD* = 35.1 ms; *N* = 160; Figure 3B). To characterise the temporal dynamics of cardiac activity relative to confidence rating scale onset, we computed trial-locked heart rate time courses for each participant. A cluster-based permutation test against zero identified three significant clusters: a brief positive deviation from −3.75 to −3.25 s before rating scale onset (peak *t* = 7.75, cluster mass = 12.74, *p* = .032), a pronounced cardiac deceleration from −2.25 to 1.75 s relative to rating scale onset (peak *t* = −24.03 at 0.25 s, cluster mass = −149.82, *p* < .001), and a rebound acceleration from 2.25 to 2.75 s (peak *t* = 9.69, cluster mass = 15.05, *p* = .025). The deceleration began before the perceptual decision, and the subsequent acceleration followed the submitted confidence rating. When split by precision, the deceleration was deeper under low than high precision, with a significant precision effect spanning −3.25 to 1.75 s relative to rating scale onset (peak *t* = −9.223 at 0.75 s, cluster mass = −74.652, *p* < .001; Figure 3C), indicating that the cardiac orienting response itself was sensitive to the precision of sensory evidence. The orienting response was also modulated by stimulus emotion, with greater deceleration after an angry face was presented (happy vs angry faces; Supplementary Figure 1). To further test whether higher-order uncertainty modulated the orienting response itself, we compared rating-scale-locked heart rate between cue-congruent and incongruent trials and stable and volatile blocks, restricting the analysis to low-precision trials. Neither contrast survived cluster-based permutation correction (Supplementary Figure 2), confirming that this cardiac response to uncertainty is specific to sensory precision.

**Figure 3:**
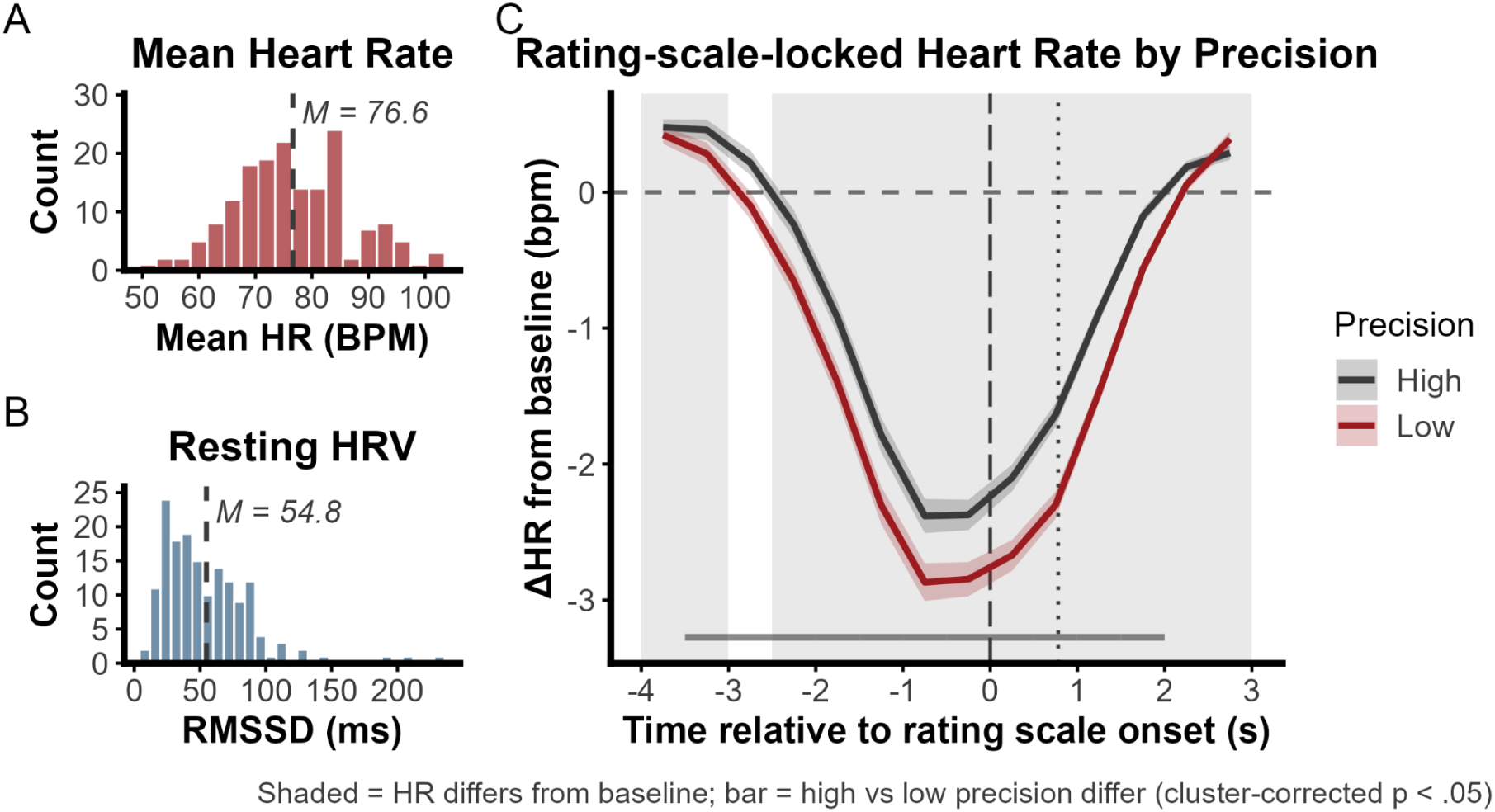
Cardiac features and rating-scale-locked heart rate by precision. A) Distribution of mean task heart rate (dashed line: *M* = 76.6 BPM). B) Distribution of resting-state HRV (*N* = 160; dashed line: *M* = 54.8 ms). C) Rating-scale-locked heart rate time course, split by sensory precision. Lines show the grand mean per condition; shaded ribbons show ± 1 SE. Grey shading indicates time bins where grand-mean heart rate deviated significantly from baseline; the horizontal bar indicates the precision-effect cluster (−3.25 to 1.75 s, *p* < .001). The dashed vertical line marks rating scale onset; the dotted line marks the median time at which the confidence rating was provided.

### Heart rate selectively predicts confidence and is modulated by sensory precision

We first tested whether trial-level heart rate predicted perceptual accuracy, to establish that any cardiac-confidence relationship could not be accounted for by general performance differences. No significant clusters survived correction (maximum observed |*t*| = 2.086, threshold *t* = 1.974; one supra-threshold cluster did not survive permutation correction, *p* = .158; BF₀₁ = 6.00; Figure 4A), indicating that trial-level cardiac activity did not reliably predict whether participants responded correctly. We then tested whether trial-level heart rate predicted confidence, controlling for accuracy and precision. A significant positive cluster was identified spanning −2.75 to 2.25 s relative to rating scale onset (peak *t* = 9.328 at 0.75 s, cluster mass = 62.950, *p* < .001; Figure 4B). Although heart rate decreased on average across this period (Figure 3C), trials with higher baseline-corrected heart rate during this window were associated with greater confidence, independently of perceptual accuracy and precision. This coupling survived when restricting to low-precision correct trials with log-transformed RT as a covariate, suggesting the result is not driven by trial difficulty (cluster −0.25 to 2.25 s, peak *t* = 5.075, *p* = .003; Supplementary Figure 3).

**Figure 4:**
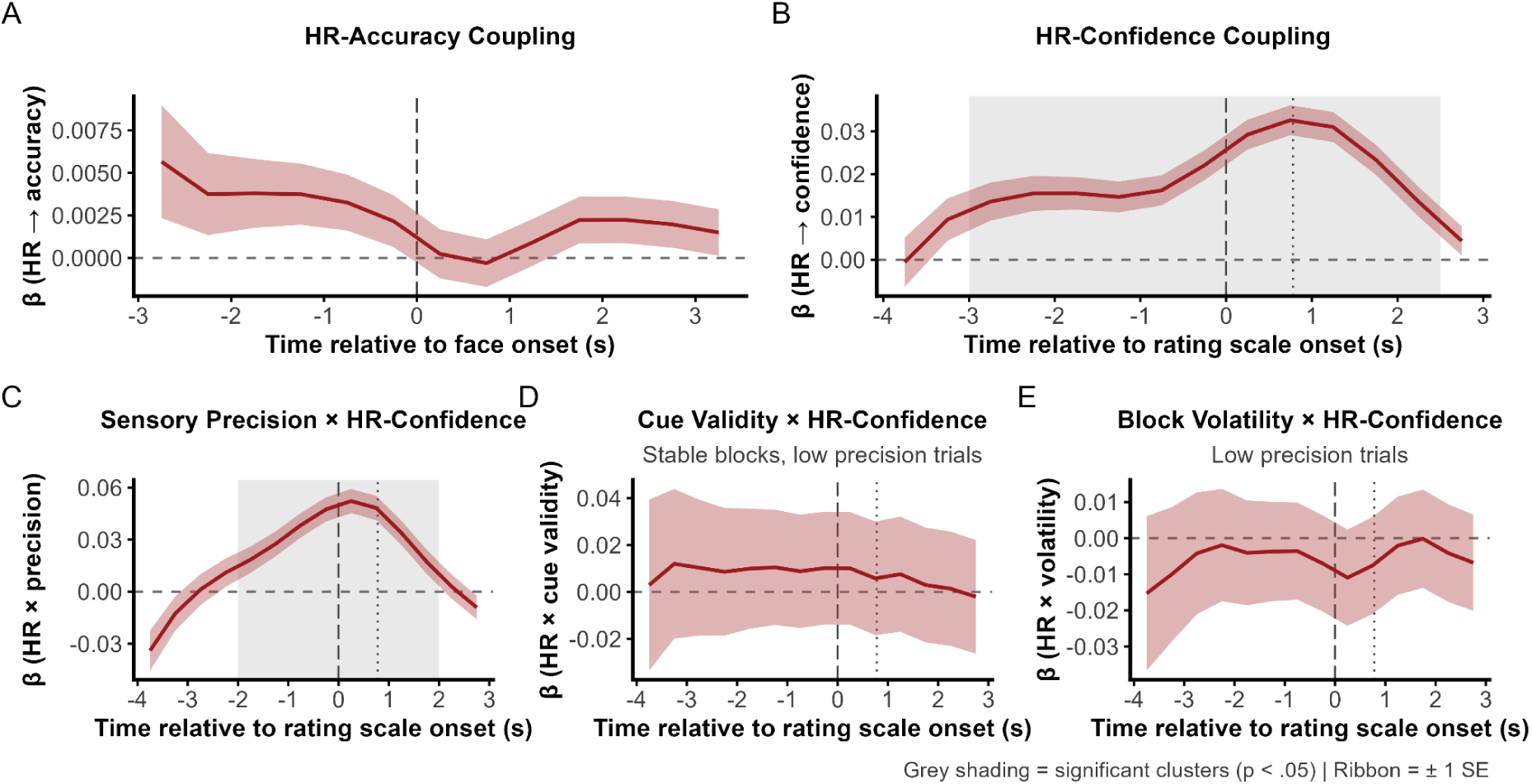
Trial-level cardiac-confidence coupling and its modulation by three sources of uncertainty. All panels show the time-resolved regression coefficient (β) from per-bin linear mixed-effects models with cluster-based permutation testing (1,000 permutations). Lines show the group-level β; shaded ribbons show ± 1 SE; grey shading indicates significant clusters (*p* < .05). A) Heart rate did not predict accuracy (face-locked; BF₀₁ = 6.00). B) Heart rate positively predicted confidence after controlling for accuracy and precision (cluster −2.75 to 2.25 s, *p* < .001). C) The heart rate × precision interaction: cardiac-confidence coupling was stronger under low than high precision (cluster −1.75 to 1.75 s, *p* < .001). D) No heart rate × cue validity interaction (low-precision, stable blocks; BF₀₁ = 5.08). E) No heart rate × block volatility interaction (low-precision, non-predictive blocks excluded; BF₀₁ = 10.24). In B-E, the dashed vertical line marks rating scale onset; the dotted line marks the median time at which the confidence rating was provided. *N* = 167.

To test whether this coupling was modulated by sensory precision, we introduced a heart rate × precision interaction term with high precision as the reference level. A significant positive cluster was identified spanning −1.75 to 1.75 s relative to rating scale onset (peak *t* = 7.492 at 0.25 s, cluster mass = 40.161, *p* < .001; Figure 4C), indicating that the coupling between heart rate and confidence was significantly stronger on low- than high-precision trials. This finding suggests that interoceptive signals contribute more to confidence formation when sensory evidence is weak. Sensory precision also modulated the mean cardiac response itself (Figure 3C), demonstrating that degraded sensory evidence affected both the cardiac response and its coupling with confidence.

To assess whether this modulation was specific to sensory precision or reflected a general sensitivity to uncertainty, we tested equivalent heart rate × uncertainty interactions for cue validity and block volatility. For cue validity, restricted to low-precision trials in stable blocks, no significant clusters were identified (maximum observed |*t*| = 0.428, threshold *t* = 1.974; BF₀₁ = 5.08; Figure 4D). For block volatility, restricted to low-precision trials with non-predictive blocks excluded, no significant clusters were identified (maximum observed |*t*| = 0.820, threshold *t* = 1.974; BF₀₁ = 10.24; Figure 4E). Bayesian analyses provided moderate evidence for the null for cue validity (BF₀₁ = 5.08) and moderate-to-strong evidence for the null for block volatility (BF₀₁ = 10.24; Supplementary Table 2). Together, these results indicate that the modulation of cardiac-confidence coupling is specific to sensory evidence precision and does not extend to the cue-validity or environmental-volatility manipulations, despite both demonstrably influencing participants’ confidence (Figure 2E, 2F).

### Cardiac signals do not predict individual differences in metacognitive efficiency

We tested whether individual differences in resting vagal tone or cardiac-confidence coupling predicted metacognitive efficiency. Group-level metacognitive efficiency, estimated by hierarchical HMeta-d on low-precision trials, was below optimal (M-ratio = 0.707, 95% HDI [0.653, 0.765]). Resting HRV did not predict metacognitive efficiency (β = 0.026 per *SD*, 95% HDI [−0.052, 0.105]; BF₀₁ = 20.61; Figure 5A, C). HR-confidence coupling strength, operationalised as the mean HR_z coefficient from per-bin OLS regressions within the −0.5 to 1.0 s window on low-precision trials (*M* = 0.0452, *SD* = 0.0818), also did not predict metacognitive efficiency (β = −0.047 per *SD*, 95% HDI [−0.126, 0.028]; BF₀₁ = 12.33; Figure 5B, D). We also computed type-2 AUROC as a non-parametric robustness check. Resting RMSSD showed no association with AUROC2 (*r*(158) = .050, *p* = .529; BF₀₁ = 4.51), consistent with the hierarchical analysis. Coupling strength was marginally associated with AUROC2 (*r*(165) = −.148, *p* = .057; BF₀₁ = 0.96 under the medium prior, 1.17 at rscale = 0.5, 1.71 at rscale = 1.0), leaving the evidence equivocal under this measure. The hierarchical M-ratio remains our primary index because it pools across participants and corrects for response bias (Fleming, 2017), but the AUROC2 result indicates the null should not be overstated.

**Figure 5:**
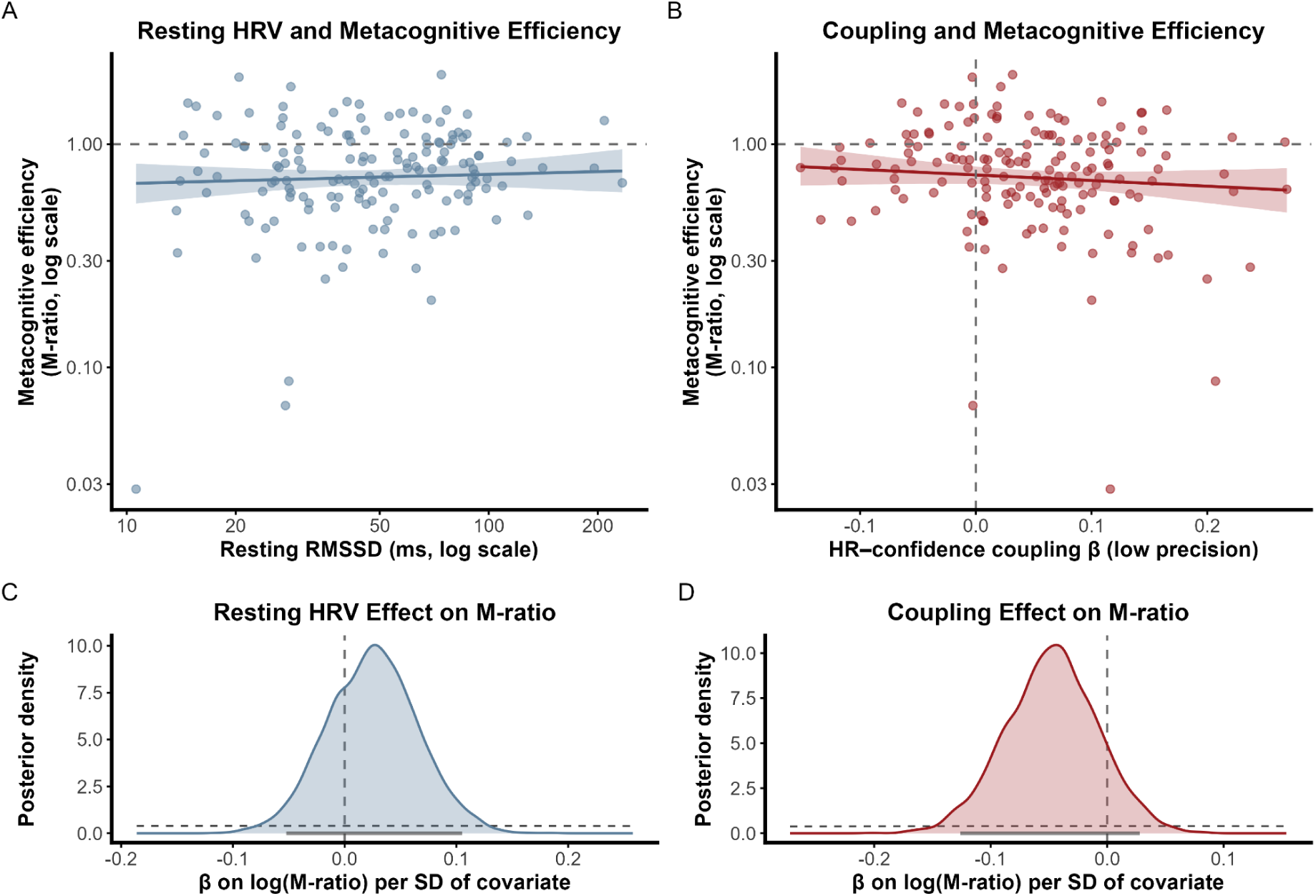
Individual cardiac metrics do not predict metacognitive efficiency. A) Resting HRV (ln-transformed) plotted against M-ratio (*N* = 160). B) HR-confidence coupling strength (mean β from the −0.5 to 1.0 s window, low-precision trials) plotted against M-ratio (*N* = 167). C) Posterior distribution of the regression coefficient for resting HRV on log(M-ratio) from the hierarchical HMeta-d model (β = 0.026, 95% HDI [−0.052, 0.105]; BF₀₁ = 20.61). D) Posterior distribution of the regression coefficient for coupling strength on log(M-ratio) (β = −0.047, 95% HDI [−0.126, 0.028]; BF₀₁ = 12.33). Shaded regions in A-B show 95% credible bands; vertical dashed lines in C- D indicate zero.

## Discussion

The aim of this study was to test whether cardiac-confidence coupling is a general response to uncertainty, or whether it is specific to the level at which sensory evidence is degraded. Three sources of uncertainty were orthogonally manipulated: sensory precision, cue validity, and block volatility. The behavioural data confirmed that all three uncertainty manipulations had the intended effect. Accuracy, reaction times, and confidence were each sensitive to precision, cue validity, and block volatility, and participants demonstrably learned the cue contingencies. Despite the universal effect on behaviour, only sensory precision significantly modulated the coupling between heart rate and confidence. This pattern is consistent with a precision-weighting account in which interoceptive signals gain influence specifically when the sensory evidence is degraded, rather than when uncertainty is introduced at higher levels of the inferential hierarchy (Barrett & Simmons, 2015).

Heart rate showed a pronounced pre-decisional deceleration that was deeper under low than high sensory precision. This finding is consistent with earlier work on the cardiac orienting response and with the hypothesis that cardiac deceleration facilitates the processing of incoming exteroceptive signals (Graham & Clifton, 1966; Porges, 2007; Skora et al., 2022). Furthermore, it suggests that the cardiac orienting response itself is sensitive to the precision of sensory evidence. The orienting response was further modulated by stimulus emotion, with a deeper deceleration following angry relative to happy faces (Supplementary Figure 1). This emotion effect did not interact with sensory precision, indicating that affective content and precision contributed to the deceleration independently. This finding is consistent with the view that stimuli signalling potential threat recruit a stronger orienting response, likely reflecting their greater motivational salience (Bradley, 2009). Whether this reflects a response to threat valence specifically or to an arousal/attention component that happens to covary with angry faces cannot be determined here. Under either interpretation, a stimulus dimension orthogonal to sensory precision shapes the pre-decisional deceleration. Temporally, the deceleration overlapped substantially with the window in which heart rate predicted confidence, raising the possibility that both phenomena reflect a shared psychophysiological process.

Heart rate positively predicted trial-level confidence, independently of accuracy and precision. This dissociates cardiac-confidence coupling from the perceptual decision it evaluates. Three accounts could explain this association. First, the coupling may reflect a bottom-up process in which cardiac afferent signals, relayed via the vagus nerve and insular cortex, are integrated into the confidence computation as a precision-weighted prior (Allen et al., 2022; Seth & Friston, 2016). Second, it could reflect a top-down process where the brain computes confidence and simultaneously drives cardiac changes, so that heart rate covaries with confidence without informing it (Critchley & Harrison, 2013). Third, the association could be a byproduct of task difficulty or attentional engagement, which could independently drive changes in both heart rate and confidence (Banellis & Cruse, 2020; Skora et al., 2022). These accounts are not mutually exclusive, and the current study design cannot distinguish directionality. However, the remaining results speak to which account is most likely.

The central finding of this study is that only sensory precision modulated the cardiac-confidence coupling. All three uncertainty manipulations affected behaviour, yet only sensory precision changed how strongly heart rate tracked confidence. The Bayes factors for the cue validity and volatility nulls support this dissociation, providing moderate evidence for the null for cue validity and strong evidence for volatility. Additionally, within low-precision trials, heart rate predicted confidence across trials matched on stimulus identity (Supplementary Figure 3), suggesting the coupling is not reducible to differences in task difficulty alone. This pattern is most consistent with a precision-weighting account in which sensory evidence determines how strongly the sensory likelihood drives the inference, so that interoceptive priors carry more weight when sensory evidence is weak (Allen et al., 2022; Feldman & Friston, 2010). We refer to this mechanism as interoceptive weighting. Cue validity and block volatility, by contrast, operate on inferential quantities rather than the sensory input itself. However, two alternatives should be considered. First, confidence ratings clustered at ceiling under high precision, limiting the variance available to detect coupling in that condition. Second, modulation of the cardiac orienting response was also specific to sensory precision (Supplementary Figure 2), leaving open whether the coupling specificity reflects an interoceptive weighting mechanism or arises because only sensory degradation elicits a cardiac response large enough to covary with confidence.

On the individual level, neither resting vagal tone nor cardiac-confidence coupling strength predicted metacognitive efficiency. Bayesian model comparison provided strong evidence for the null for both resting HRV and coupling strength when metacognitive efficiency was estimated hierarchically. Under narrower priors and when indexed by AUROC2, the evidence was weaker, suggesting that the conclusion depends in part on how metacognitive efficiency is operationalised. Still, these results suggest the interoceptive weighting of cardiac signals in confidence computations is a within-trial dynamic rather than a stable individual-differences trait. However, robust trial-level effects in cognitive tasks do not always produce reliable individual differences (Hedge et al., 2018). In the present study, this may have been compounded by the fact that resting HRV was derived from a separate MRI session rather than during the task session. Still, the trial-level association and the individual-difference nulls converge on a common interpretation: the cardiac signal may be associated with higher confidence without improving metacognitive efficiency (Allen et al., 2016). Whether the association between cardiac signals and higher confidence is adaptive (for instance, by facilitating decisional commitment under uncertainty; Desender et al., 2019) or maladaptive (by inflating confidence beyond what the evidence supports; Zylberberg et al., 2016) remains an open question. Future work could address this directly by testing whether cardiac-confidence coupling differs between correct and incorrect trials: a coupling that selectively amplifies confidence on correct trials would be consistent with an adaptive signal. A complementary approach would be to incorporate cardiac signals into computational models of evidence accumulation and metacognition to specify how they enter the confidence computation.

Several limitations of the current study should be considered. First, the current study is correlational with respect to cardiac signals: heart rate was measured but not manipulated. Establishing directionality will require experimental manipulation of cardiac signals independently of the decision process (Garfinkel & Critchley, 2016), for instance through pharmacological modulation of cardiac activity (Tyrer et al., 2025) or non-invasive cardiac stimulation. Second, confidence ratings under high sensory precision clustered at ceiling, limiting the variance available in this condition. Whether the stronger coupling under low precision reflects a genuine increase in interoceptive weighting or partly the scale’s greater sensitivity in that condition cannot be determined from the current design. Future work should systematically vary task difficulty to prevent ceiling confidence under high precision, allowing the interaction to be tested with comparable variance across conditions. This ceiling also necessitated restricting the cue validity and volatility analyses to low-precision trials. Third, the behavioural effects of sensory precision were substantially larger than those of cue validity and volatility. Consequently, we cannot rule out that a stronger manipulation of higher-order uncertainty would also reveal modulation of the coupling at those levels. Fourth, the present findings are specific to face-emotion discrimination, a task domain that may engage affective processing in addition to perceptual decision-making (Garfinkel et al., 2014). Because every stimulus carried emotional content, the precision manipulation cannot dissociate degradation of the sensory likelihood as a whole from degradation of its emotional component; a non-emotional control such as the face-versus-house task with superimposed white visual noise used by Lawson et al. (2021) would be needed to resolve this.

Confidence is often treated as though it were computed from the evidence that informed the decision, with added noise (Fleming, 2024). Our findings here challenge this view, showing that the covariation between cardiac signals and confidence is temporally organised and specific to the level of the inferential hierarchy at which sensory evidence is degraded. At the same time, this coupling does not appear to improve metacognitive accuracy: it is associated with higher confidence but not with higher metacognitive efficiency. This dissociation matters clinically. Alterations in interoceptive processing have been reported in anxiety, depression, and psychosis (Brewer et al., 2021), conditions in which metacognition is also often disrupted (Hoven et al., 2019; Rouault et al., 2018). Yet, a mechanistic account connecting interoceptive dysfunction to disrupted metacognition is lacking. The present findings offer a starting point: if the cardiac-confidence association is strongest when sensory evidence is degraded, then disrupted interoceptive processing should produce its largest effects on confidence specifically under those conditions. Establishing whether this association is causal, and identifying the neural circuitry that mediates it, are the necessary next steps toward a mechanistic account of how signals from the body shape how we evaluate our own decisions.

## Declaration of generative AI and AI-assisted technologies in the manuscript preparation process

During the preparation of this work the authors used Claude to assist with writing, editing, and coding tasks. After using this tool, the authors reviewed and edited the content as needed and take full responsibility for the content of the published article.

## Acknowledgements

This work was carried out at the Center of Functionally Integrative Neuroscience, Aarhus University.

## Supplementary Material

**Supplementary Table 1:**
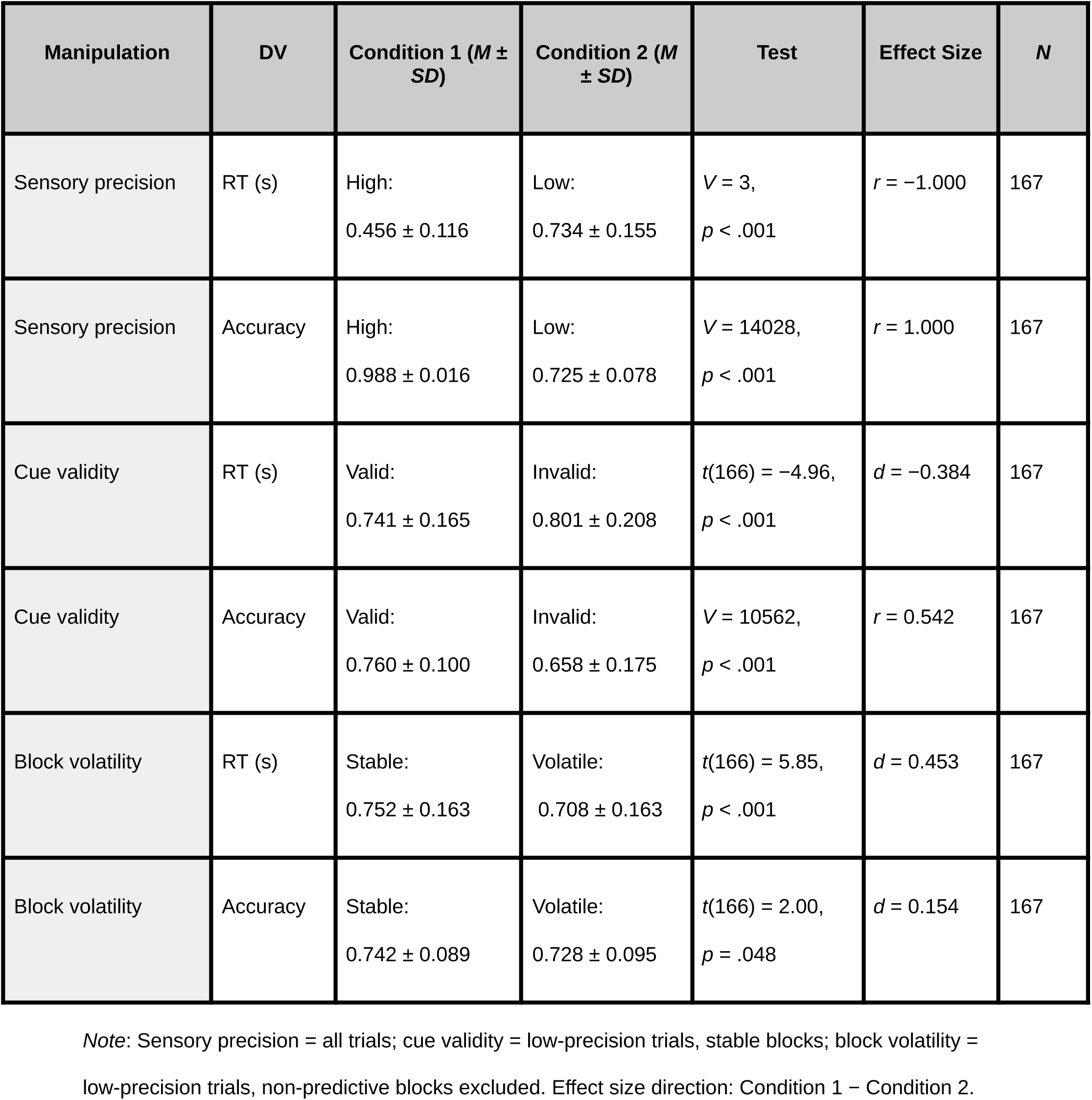
Behavioural performance across uncertainty manipulations.

**Supplementary Table 2:** Bayes factor sensitivity analysis for null cardiac-coupling results.

| **Analysis** | **Subset** | **Window (bins)** | **Mean β (SD)** | ***t*(166)** | ***p*** | ***r* = 0.5** | ***r* = √2/2** | ***r* = 1** | **Evidence** |
|---|---|---|---|---|---|---|---|---|---|
| HR → accuracy (4A) | All trials | 0.0-3.5 s (7) | 0.0014 (0.0160) | 1.16 | .248 | 4.35 | 6.00 | 8.38 | Moderate |
| HR × cue validity (4D) | Low prec., stable | −0.5-1.0 s (3) | 0.0418 (0.4161) | 1.30 | .196 | 3.70 | 5.08 | 7.08 | Moderate |
| HR × volatility (4E) | Low prec., predictive | −0.5-1.0 s (3) | −0.0070 (0.1811) | −0.50 | .616 | 7.34 | 10.24 | 14.37 | Moderate/ strong |

*Note.* BF<sub>01</sub> = Bayes factor in favour of H<sub>0</sub>, computed via JZS one-sample *t*-test at three prior widths (*r* = Cauchy scale parameter; Morey et al., 2026). BF<sub>01</sub> > 3 indicates moderate evidence for the null; BF<sub>01</sub> > 10 indicates strong evidence. All three analyses support the null at every prior width. The evidence category for block volatility shifts from strong (*r* = √2/2) to moderate (*r* = 0.5), but the qualitative conclusion is unchanged.

**Supplementary Figure 1:**
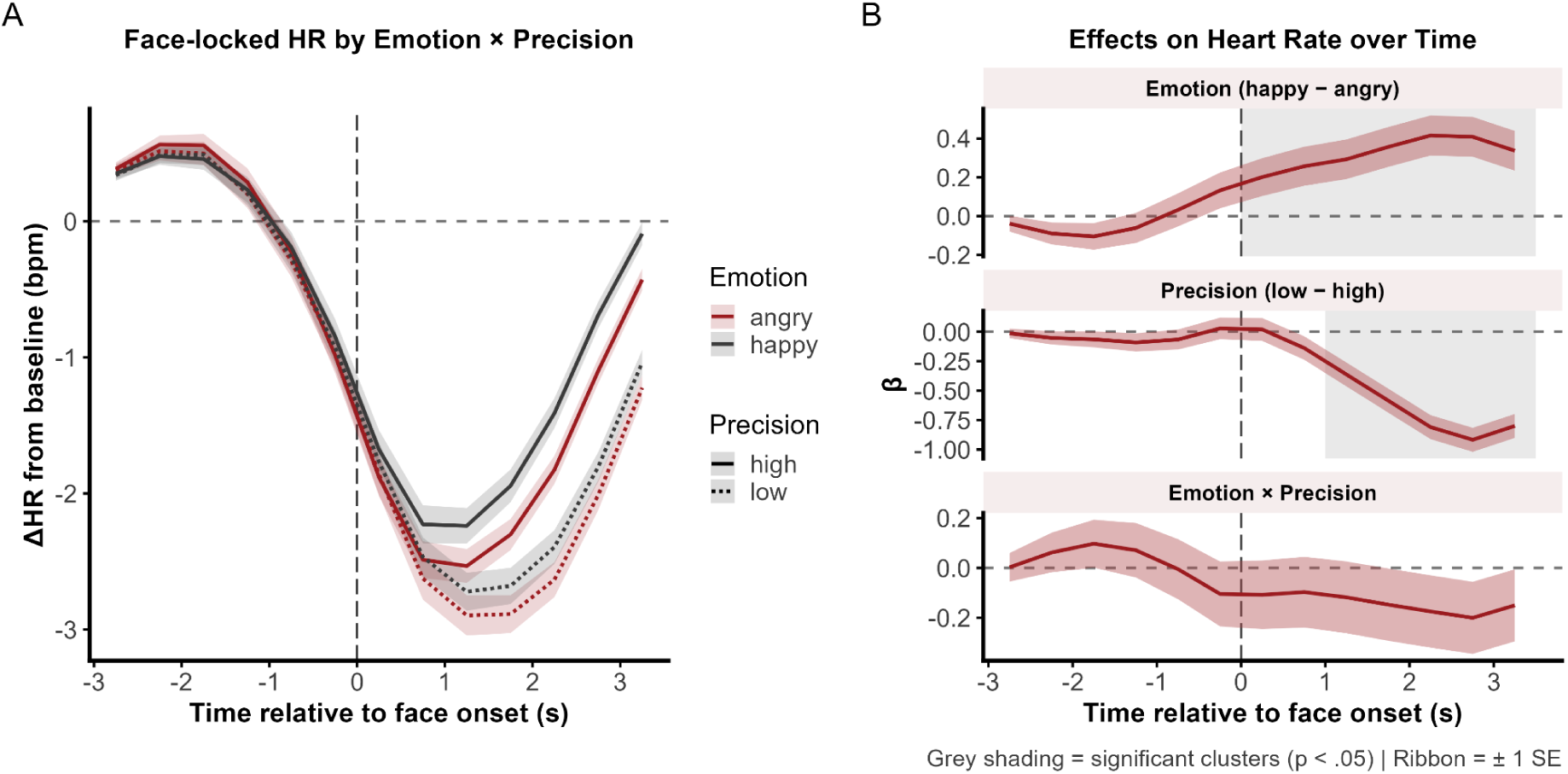
Face-locked heart rate by stimulus emotion and sensory precision. Both panels show time-resolved analyses of heart rate in the face-locked epoch; shaded ribbons show ± 1 SE; grey shading indicates significant clusters (*p* < .05). A) Grand-mean heart rate time courses split by stimulus emotion (angry vs. happy) and sensory precision (high vs. low). B) Time-resolved regression coefficients (β) from per-bin linear mixed-effects models with cluster-based permutation testing for the emotion main effect, precision main effect, and their interaction. A significant cluster for the emotion main effect (happy vs. angry) was observed over [0.25, 3.25] s (peak *t* = 4.042, *p* = .026). A significant cluster for the precision main effect was observed over [1.25, 3.25] s (peak *t* = −9.034, *p* = .002). The emotion × precision interaction was not significant (maximum observed |*t*| = 1.386). *N* = 167.

**Supplementary Figure 2:**
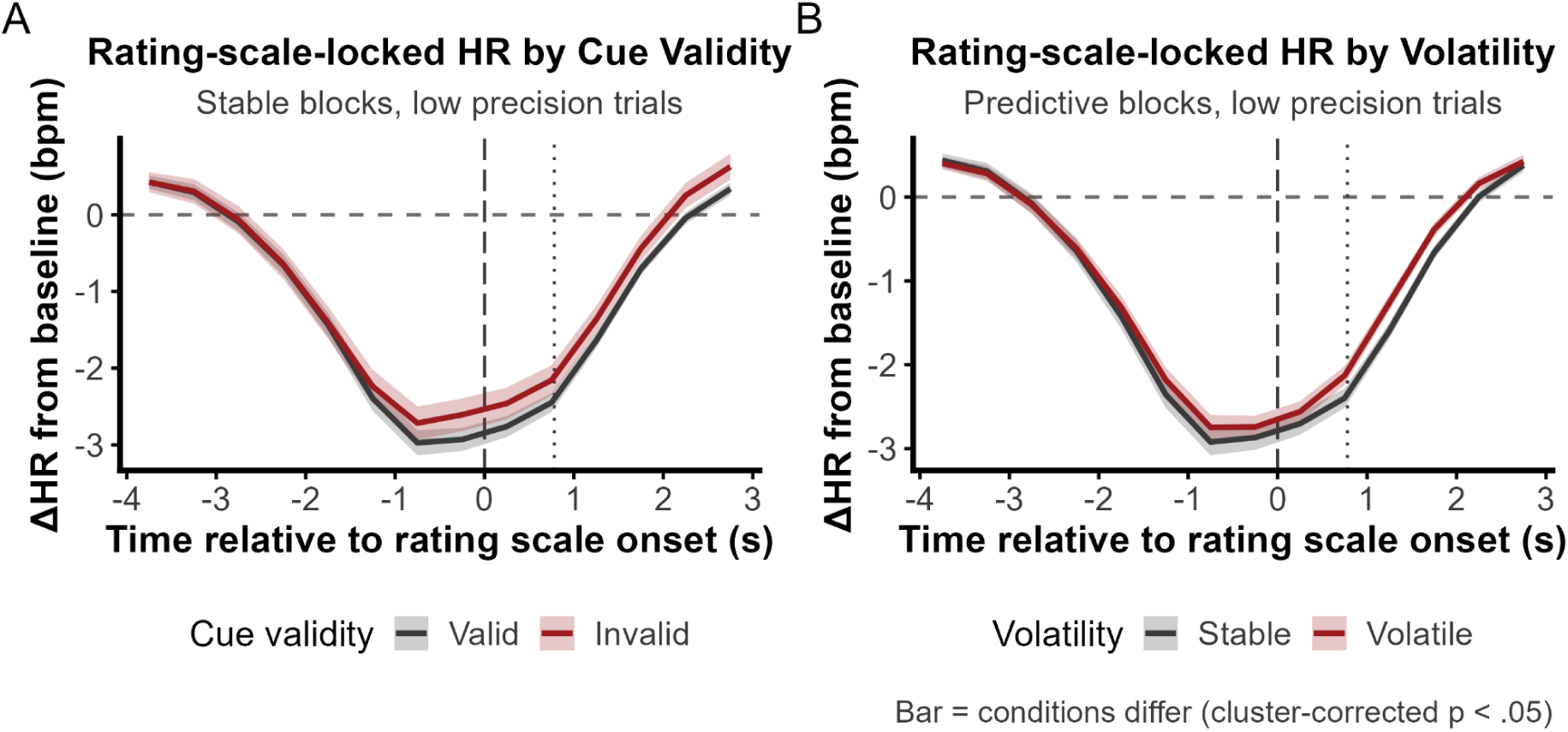
Rating-scale-locked heart rate by cue validity and block volatility. Rating-locked heart rate time courses for low-precision trials, baseline-corrected to the 1 s preceding cue onset. Lines show the grand mean per condition; shaded ribbons show ± 1 SE. A) Cue congruent versus cue incongruent trials (stable blocks only). No clusters survived cluster-based permutation correction (largest cluster: [−0.25] s, mass = 2.05, peak *t* = 2.045 at −0.25 s, *p* = .198; 1000 permutations). B) Stable versus volatile blocks (non-predictive blocks excluded). No clusters survived cluster-based permutation correction (largest cluster: [0.75, 1.75] s, mass = 8.42, peak *t* = 3.159 at 1.25 s, *p* = .074; 1000 permutations). Neither cue validity nor block volatility modulated the rating-locked orienting response, indicating that this cardiac signature is specific to sensory precision.

**Supplementary Figure 3:**
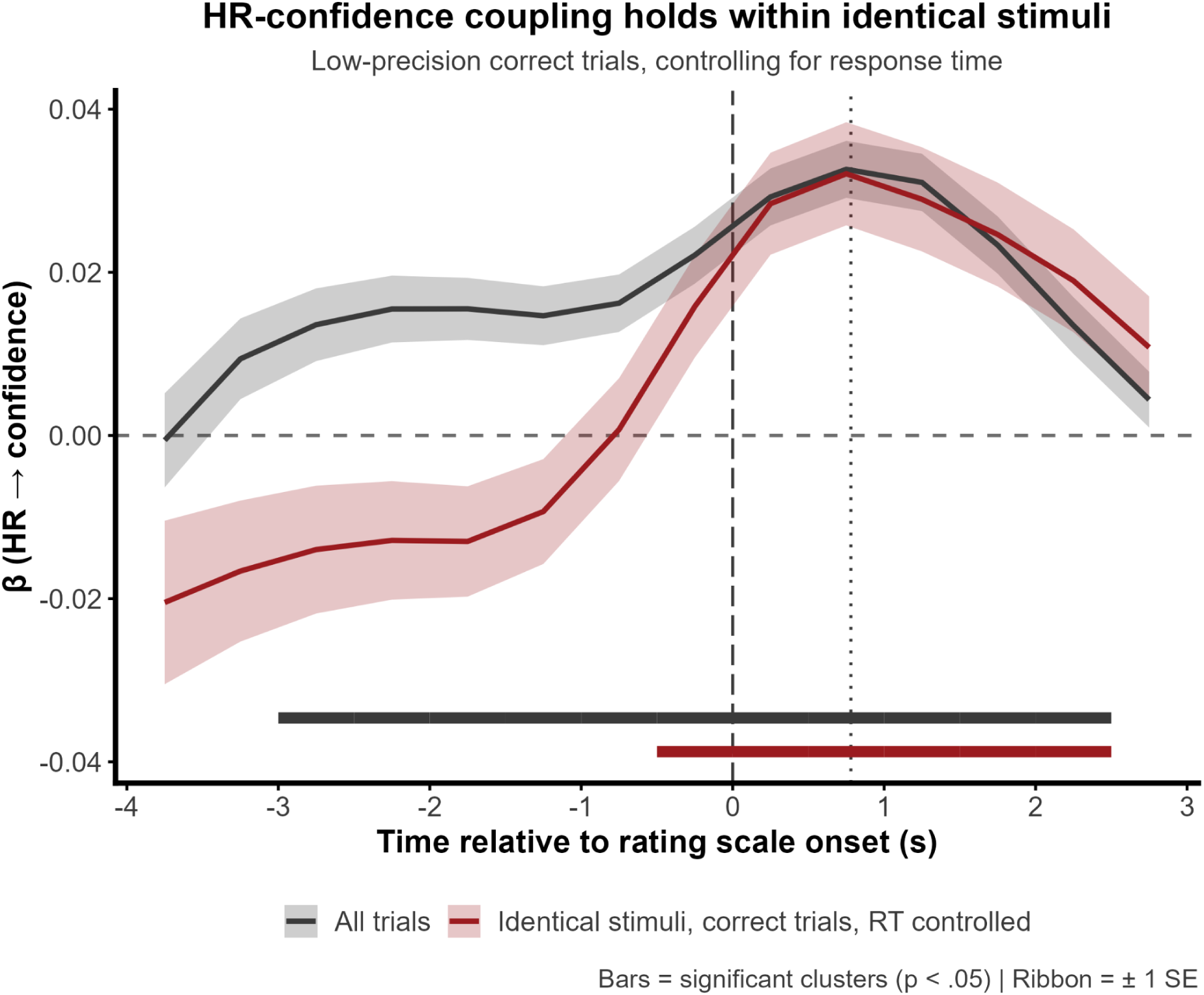
HR-confidence coupling holds within identical stimuli and after controlling for response time. Grey: original coupling analysis across all trials (confidence_z ∼ HR_z + accuracy + precision + (1|sID); reproduces Figure 4B). Red: restricted to low-precision correct trials with log-transformed RT as a covariate (confidence_z ∼ HR_z + logRT_z + (1|sID/face_outcome)), holding stimulus difficulty, perceptual outcome, and response speed constant. Horizontal bars indicate significant clusters (*p* < .05; all trials: −2.75 to 2.25 s, *p* < .001; identical stimuli: −0.25 to 2.25 s, *p* = .003). The coupling emerges later and is attenuated in the controlled analysis, but survives from rating onset onward, peaking at 0.75 s. The dashed vertical line marks rating-scale onset; the dotted line marks the median time at which the confidence rating was provided. Ribbons show ± 1 SE. *N* = 167.

